# A neural mechanism of cognitive reserve: The case of bilingualism

**DOI:** 10.1101/2022.09.20.508678

**Authors:** W. Dale Stevens, Naail Khan, John A. E. Anderson, Cheryl L. Grady, Ellen Bialystok

## Abstract

Cognitive Reserve (CR) refers to the preservation of cognitive function in the face of age-or disease-related neuroanatomical decline. While bilingualism is known to contribute to CR, the extent to which, and what particular aspect of, second language experience contributes to CR are debated, and the underlying neural mechanism(s) unknown. Intrinsic functional connectivity reflects experience-dependent neuroplasticity that occurs across timescales ranging from minutes to decades, and may be a neural mechanism underlying CR. To test this hypothesis, we used voxel-based morphometry and resting-state functional connectivity analyses of MRI data to compare structural and functional brain integrity between bilingual and monolingual older adults, matched on cognitive performance using a rigorous propensity score matching technique, and across levels of second language proficiency measured as a continuous variable. Bilingualism, and degree of second language proficiency in particular, were associated with lower grey matter integrity in a hub of the default mode network – a region that is particularly vulnerable to decline in aging and dementia – but preserved functional network organization that resembled the young adult brain. Our findings confirm that lifelong bilingualism contributes to CR through experience-dependent maintenance of optimal functional network structure of the domain-general attentional control network across the lifespan.

---

Multiple factors contribute to Cognitive Reserve (CR) across the lifespan, such as education, occupation, and intellectual and social engagement (Cabeza et al., 2018; Song et al., 2022; Stern, 2021). There is compelling evidence that lifelong bilingualism also contributes to CR (Anderson et al., 2020; Bialystok, 2021). Relative to their monolingual peers, bilingual older adults 1) have better cognitive function, given similar levels of neuroanatomical decline, 2) demonstrate signs of dementia later in life, 3) show more significant neuroanatomical pathology, given similar levels of dementia, and 4) show more rapid rates of cognitive decline in later stages of dementia (Bialystok, 2021). Bilingual individuals must continuously exert cognitive control to appropriately direct their attention towards one situationally relevant language of two that are simultaneously active in mind and competing for attention (Kroll et al., 2014; van Heuven et al., 2008). Evidence suggests that this ongoing need for selection leads to an adaptation in domain-general attentional control that enhances multiple subordinate cognitive control process across a range of tasks and mitigates typical age-related cognitive decline (Bialystok, 2015; Bialystok and Craik, 2022). This adaptation is the foundation for CR in bilingualism (Bialystok, 2021). Thus, bilingualism serves as a useful proxy for investigating neural mechanisms underlying CR. Given the evidence that lifelong bilingualism is a CR factor, two predictions are that 1) given equal neuroanatomical health, bilinguals will have better cognitive performance than monolinguals and 2) given equal cognitive performance, bilinguals will have poorer neuroanatomical health than monolinguals. Both relationships have been confirmed in recent work (Anderson et al., 2021; Berkes et al., 2021; Brian T. Gold et al., 2013; Perani et al., 2017; Schweizer et al., 2012).

The neural mechanism underlying CR in bilingualism is unknown. Studies investigating differences in brain structure associated with bilingualism have produced mixed, often contradictory results. For example, while some studies have shown better structural integrity in bilingual than monolingual individuals (John A.E. Anderson et al., 2018; Luk et al., 2011; Marin-Marin et al., 2020; Olsen et al., 2015), others have shown the opposite (Anderson et al., 2021; Brian T. Gold et al., 2013; Schweizer et al., 2012), or dissociable patterns of grey vs. white matter integrity (Macbeth et al., 2021). While the underlying neural mechanisms by which augmented cognitive engagement across the lifespan leads to preservation of cognitive faculties in later life are undetermined, there might be multiple pathways to CR (Oh et al., 2018). Another possibility is that CR might result from adaptive shifts in cognitive strategy or more efficient task-related engagement of brain regions or circuits (Abutalebi et al., 2012; Anderson et al., 2021; B. T. Gold et al., 2013). A third possibility is that experience-dependent maintenance or enhancement of the brain’s long-term intrinsic functional network integrity could be a neural mechanism underlying CR (Chan et al., 2021, 2018; N. Franzmeier et al., 2017; Nicolai Franzmeier et al., 2017b; Weiler et al., 2018; Wig, 2017).

RSFC, which can be quantified by analyzing spontaneous fluctuations of the blood-oxygen-level-dependent (BOLD) signal in resting-state functional MRI (fMRI) data, reflects the brain’s intrinsic large-scale functional network architecture [for example, see refs. (Biswal et al., 1995; Fox and Raichle, 2007)]. RSFC plays a central role in human cognition through a Hebbian-like mechanism, whereby repeated experience-driven coupling or decoupling of task-related activity among brain regions over time lead to enduring increases or decreases in RSFC among these regions, respectively (Stevens and Spreng, 2014). This, in turn, facilitates or inhibits future coupling (Kelly and Castellanos, 2014), thus enhancing or optimizing cognitive performance with experience over time. Thus, RSFC reflects a previously unappreciated mechanism of long-term neurocognitive plasticity that allows past experiences to mold sustained functional network architecture to facilitate future cognitive performance. Research on RSFC has demonstrated that 1) it can be modulated by ongoing or recent experience across functionally relevant brain regions/circuits/networks on relatively short timescales ranging from minutes to days (Martin et al., 2021); 2) the degree of sustained experience-dependent change predicts learning, memory consolidation, and future performance (Lewis et al., 2009; Stevens et al., 2010; Tambini et al., 2010); 3) patterns of RSFC are correlated with individual differences in personality, behavior, and cognition (Adelstein et al., 2011; Nostro et al., 2018); and 4) differential patterns of RSFC are associated with neurodevelopmental and neurodegenerative disorders, and the extent of neurocognitive decline/preservation in old age (Stevens and Spreng, 2014; Wig, 2017). Thus, RSFC reflects a critical process underlying long-term neurocognitive plasticity that facilitates the development, specialization, stabilization, and preservation of large-scale functional brain networks across the lifespan. While there may be multiple pathways to CR, we propose that experience-dependent maintenance of intrinsic functional networks is a fundamental neural mechanism that could underly the phenomenon of CR in lifelong bilingualism.

Various RSFC analytic techniques can be used to parcellate the brain into multiple functionally dissociable regions that are organized into large-scale functional sub-networks with varying degrees of spatial resolution (Power et al., 2011; Schaefer et al., 2018; Wig et al., 2014; Yeo et al., 2011). Previous work has focussed on three specific sub-networks that play foundational roles in higher cognitive functioning (Grady et al., 2016; Setton et al., 2022; Spreng et al., 2013; Spreng and Schacter, 2012): 1) the dorsal attention network (DAN), involved in externally focussed cognition; 2) the default mode network (DMN), involved in internally focussed cognition; and 3) the frontoparietal control network (FPCN), which facilitates volitional control over the locus of attention by flexibly alternating its coupling between the DAN vs. DMN during externally vs. internally focussed cognition, respectively (Spreng et al., 2010). Critically, different nodes within these networks play specialized roles in facilitating within-network communication and cross-network interactions among these networks, a set of dynamics that can be characterized and quantified using graph theory analyses (Rubinov and Sporns, 2010; Wig et al., 2011). Previous work has identified two discrete nodes of the FPCN within the left middle frontal gyrus (MFG) that play opposite but complementary roles – an anterior region (aMFG; Brodmann area (BA) 9) that has dense within-network connections but little cross-network connectivity with other networks (i.e., a “provincial hub node”), and a posterior region (pMFG; BA 6), which has high betweenness centrality, a measure of the extent to which a node contributes to communication across networks (i.e., a “connector node”) (Spreng et al., 2013). These two nodes are thought to play critical roles in attentional control through mediating within-network communication of the FPCN, and dynamic flexibility of cross-network coupling with the DAN and DMN, respectively (Spreng et al., 2013). There is evidence that age-related cognitive decline is related to functional connectivity changes most prominently within the FPCN (Campbell et al., 2012), and of the MFG in particular (Spreng et al., 2018).

While some aspects of cognition decline in old age, such as working memory, episodic memory, and processing speed, others remain stable or improve across the lifespan, such as semantic memory and verbal knowledge (Park et al., 2002). This pattern of age-related change has been characterized as a shift from fluid to crystalized cognition in later life (Craik and Bialystok, 2006), and it is correlated with increased RSFC of the left middle frontal gyrus, a core node of the FPCN, with the DMN – a phenomenon characterized by the “default-executive coupling hypothesis of aging” (DECHA) (Spreng and Turner, 2019). The DECHA is consistent with prior work demonstrating that 1) unlike young adults, older adults “fail to deactivate” the DMN during externally focussed cognition (Grady et al., 2006; Lustig et al., 2003; Miller et al., 2008) and 2) age-related differences in how the FPCN connects with other networks impacts both the function of these other networks and cognitive ability in older adults (Grady et al., 2016).

Here, we investigated the hypothesis that lifelong bilingualism contributes to CR by mitigating the typical age-related changes in functional connectivity of the FPCN described above (e.g., DECHA (Spreng et al., 2018)), specifically, by bolstering functional integration within the FPCN and maintaining the dynamic flexibility of its cross-network coupling with the DMN and DAN (Spreng et al., 2010). To isolate the extent to which lifelong bilingualism contributes to CR, one must either compare cognitive performance between monolingual and bilingual groups equated for neuroanatomical health (Berkes et al., 2021), with the expectation that bilinguals will outperform monolinguals; or alternatively, compare differences in brain structure and function between groups equated for cognitive performance (John A.E. Anderson et al., 2018), with the expectation that bilinguals will show more prominent signs of neuroanatomical aging. Here, we took the latter approach, and compared brain structure and intrinsic functional network integrity in groups of monolingual vs. bilingual older adults equated on multiple measures of cognitive performance and demographic variables using a rigorous propensity score matching (PSM) technique. Recent research has also supported the view that bilingualism is better characterized and investigated as a multidimensional set of continuous variables on which individuals vary based on language experience, rather than a dichotomous variable separating individuals into ostensibly homogenous groups (Dash et al., 2022; DeLuca et al., 2019; Gullifer and Titone, 2020; Macbeth et al., 2021; Sulpizio et al., 2020). Moreover, previous behavioural research has implicated the active use and proficiency of a second language as the critical factor that might contribute most to CR in bilingual older adults (Calabria et al., 2020). Thus, we also assessed the degree to which these measures of structural and functional brain integrity are associated with second language proficiency, measured as a continuous variable across all older adults in our sample. We predicted that bilingual older adults would show more prominent signs of neuroanatomical aging (i.e., “older looking” brain structure), but preserved integrity of intrinsic functional network connectivity (i.e., “younger looking” brain functional organization) relative to cognitively matched monolinguals, who would show typical patterns of degraded functional network connectivity associated with aging. Specifically, we hypothesized that 1) bilingual older adults will show lower grey matter integrity in brain regions that are particularly susceptible to age-related decline (e.g., DMN regions); 2) bilinguals will show less default-executive coupling, i.e., RSFC between the MFG and the DMN, a hallmark characteristic of neurocognitive aging (Spreng and Turner, 2019); 3) bilinguals will show preserved flexibility of cross-network RSFC of the MFG with the DAN and DMN; and 4) these measures of structural and functional integrity will be associated with second language proficiency specifically, across all older adults.

## Results

Ninety-three right-handed older adult participants (age: mean ± SD = 74.04 ± 3.86, range = 68-85; 63 female) with no history of heart disease, neurological or psychological disorders, or traumatic brain injuries, were recruited from the local area surrounding York University, Toronto, Canada (Note: An additional 3 participants were recruited but not included in this study, due to excessive head motion in the MRI scanner). Behavioral and structural MRI data from a subset of these participants have been analyzed in previous reports (Anderson et al., 2021, 2017; John A.E. Anderson et al., 2018). The Language and Social Background Questionnaire [LSBQ (John A. E. Anderson et al., 2018)] was used to categorize participants as monolingual or bilingual (LSBQ composite score) and to assign a score on second language proficiency (LSBQ factor), a continuous variable. Participants were interviewed via telephone to validate language status, and those who could not be reliably identified as monolingual or bilingual were not included in the study. Forty-seven participants (29 female) were identified as bilingual and 46 (34 female) were identified as monolingual. All participants completed the D-KEFS battery (Delis et al., 2001), comprising multiple neuropsychological tests of executive function, and the Mini-Mental State Examination [MMSE (Folstein et al., 1975)]. We used a PSM technique to equate the groups across performance on multiple cognitive measures (MMSE, Trail Making Task (letter-number-switching score), Shipley-2 verbal score, Shipley-2 block patterns score), as well as demographic variables (age, education, gender), similar to previous work (John A.E. Anderson et al., 2018). We used voxel-based morphometry (VBM) analysis of structural MRI data, and RSFC analyses of resting-state fMRI data, to compare neuroanatomical and intrinsic functional network integrity, respectively, between the groups and across second language proficiency.

### Propensity score matching on cognitive measures and demographic variables

Previous research has used sequential bivariate matching; however, this fails to account for potential multivariate interactions between the matching variables and may introduce bias into participant selection when using a high dimensional set of matching criteria (John A.E. Anderson et al., 2018; Ho et al., 2007; Rosenbaum and Rubin, 1983). With PSM, a propensity score for each individual is obtained by using logistic regression to predict group membership, given a set of observed covariates. Participants from one group are then individually matched to those in the second group based on propensity scores (Rosenbaum and Rubin, 1983). PSM can account for multivariate interactions, minimizes selection error when using a large number of matching variables, and is better suited for smaller sample sizes, as is often the case in neuroimaging studies (Austin and Steyerberg, 2015; Ho et al., 2007; Rosenbaum and Rubin, 1983). To prioritize matching the monolingual and bilingual groups on cognitive performance, PSM was done using the MatchIt R package (Ho et al., 2011) in a 2-stage hierarchical manner. The first stage matched the groups on 4 neuropsychological measures: MMSE, Trail Making Task (letter-number-switching score), Shipley-2 verbal score, and Shipley-2 block patterns score. Using the propensity scores, each bilingual participant was matched to a monolingual; bilinguals that could not be matched, and any remaining monolinguals, were removed from the matched-group analyses. The distribution of propensity scores for the matched monolingual and bilingual groups, as well as for the excluded participants from each of the unmatched groups, are shown in Supplementary Figure 1a. Once the groups were matched on cognitive performance, they were then matched on 3 demographic variables (age, education, and sex) using the same procedure. The distribution of propensity scores for the two matched groups following the second stage are shown in Supplementary Figure 1b. The absolute standardized mean difference between monolinguals and bilinguals on each of the matching variables for the matched vs. unmatched groups are show in Supplementary Figure 1c. Following the PSM procedure, the final group sizes were 39 monolinguals (age: mean ± SD = 73.51 ± 3.25, range = 68-79; 30 female) and 39 bilinguals (age: mean ± SD = 73.87 ± 4.00, range = 69-83; 23 female). The matched groups did not differ significantly on any of the cognitive or demographic variables (Table 1).

**Table 1.**
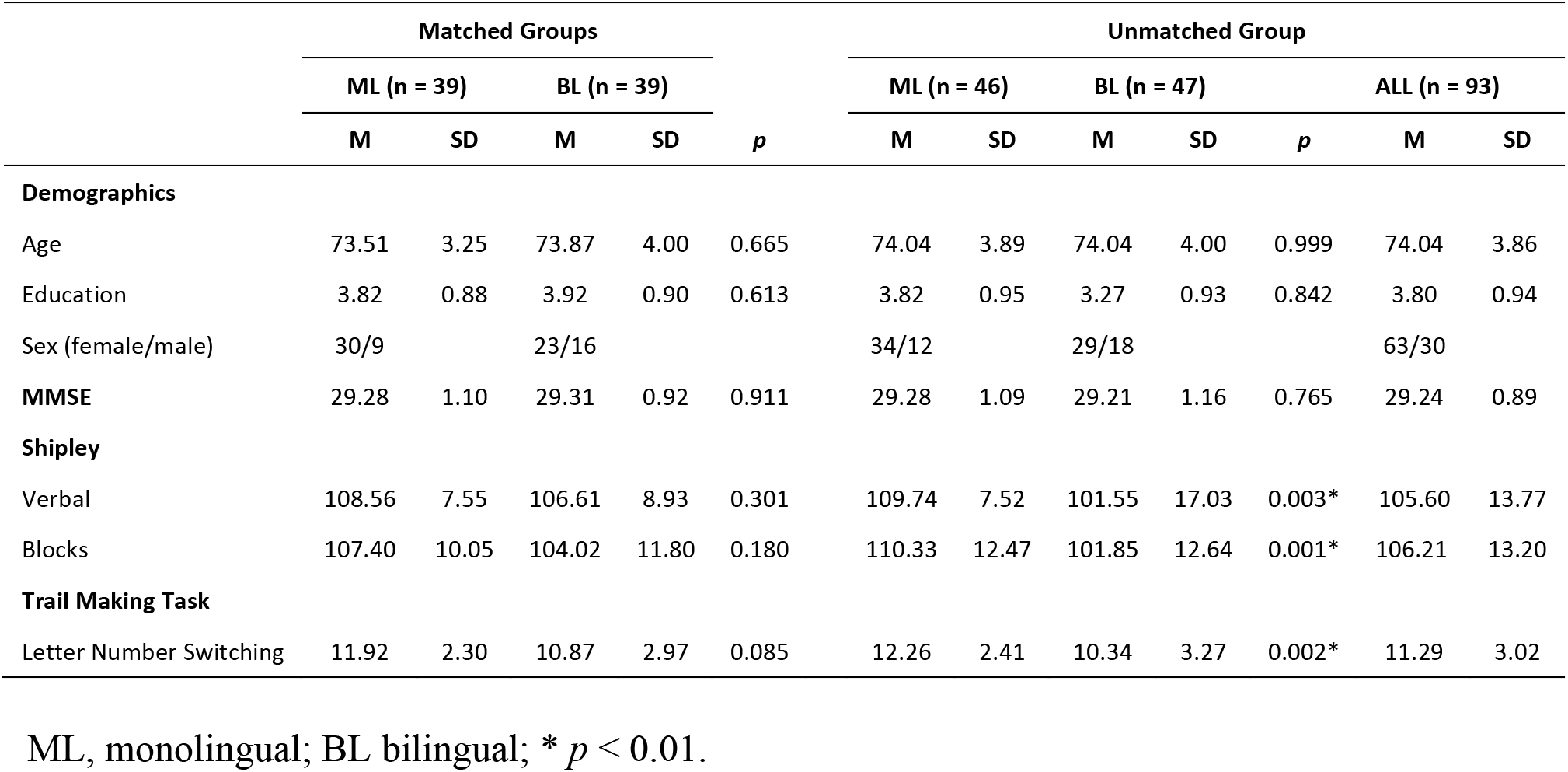
Participant information, demographics, and test scores.

### Second language proficiency as a continuous variable across all older adults

The quantity and quality of exposure to multiple languages is extremely variable across individuals, and most have at least some experience with a second language. A recent study reported that more than 80% of nearly 1,000 respondents in the United Kingdom who claimed to be monolingual English-speakers had learned other languages at some point in their lives, sometimes to high degrees of fluency (Castro et al., 2022). Previous behavioural research has implicated the active use and proficiency of a second language as the critical factor that might contribute most to CR in bilingual older adults (Calabria et al., 2020). In addition to the composite score, the LSBQ quantifies second language proficiency based on a factor analysis, which produced a latent variable comprising clustered items that were identified as representing second language proficiency. The eigenvalue from this latent variable is used as the quantitative measure of second language proficiency (for details, see ref. 59). Thus, an important subscale of the LSBQ is second language proficiency, on which individual participants vary along a continuum (John A. E. Anderson et al., 2018). To test the hypothesis that second language proficiency is a key factor that contributes to CR in bilingualism, we correlated normalized scores on this factor with normalized measures of neuroanatomical and intrinsic functional network integrity across our entire sample of 93 older adults included in the study.

### Monolingual adults have higher grey matter density in a core hub of the DMN

To test the hypothesis that monolingual older adults will have higher neuroanatomical integrity than bilinguals when the groups are equated on cognitive performance, we quantified grey matter density across the brain using VBM analysis (FSL-VBM software suite) of T1-weighted structural MRI images. This analysis identified a large cluster in the posterior cingulate cortex (pCC), a core hub of the DMN (Buckner et al., 2009; Hagmann et al., 2008; van Oort et al., 2014), that showed higher grey matter density in monolingual older adults relative to the bilinguals [p < 0.05, cluster corrected for multiple comparisons using ETAC (Cox, 2019)] (Fig. 1a-b). Morever, grey matter density in the pCC was negatively correlated with second language proficiency across the entire sampe of older adults (r = −0.30; p = 0.002, 2-tailed; Fig. 1c).

**Figure 1.**
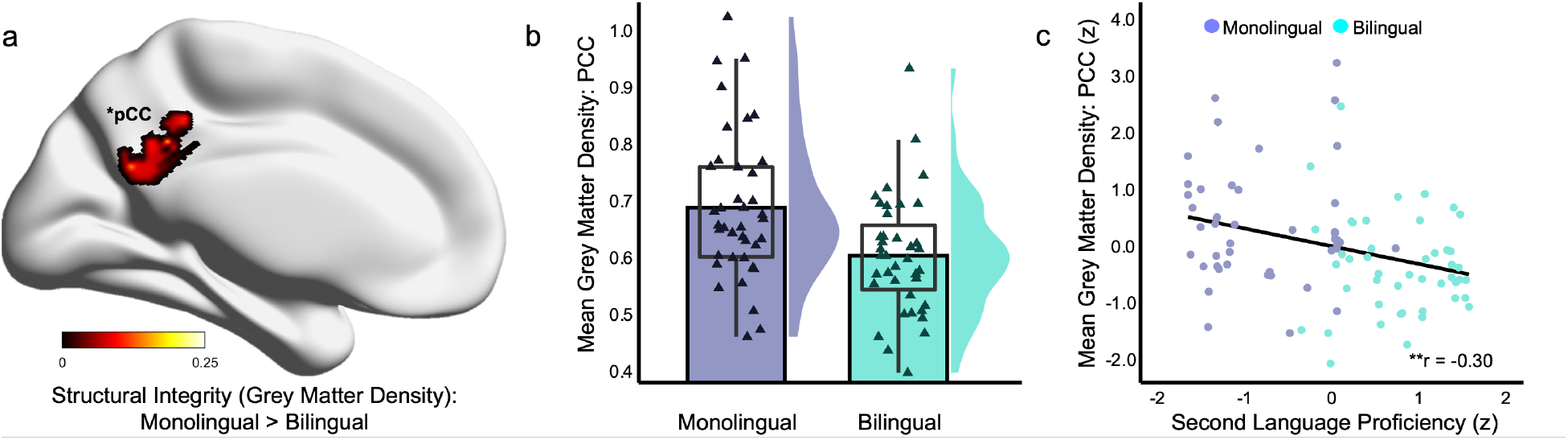
Differences in structural integrity estimated using VBM. (a) Comparing groups matched on cognitive performance using PSM, monolinguals have higher grey matter density than bilinguals in the posterior cingulate cortex, a hub of the DMN (p < 0.05, ETAC corrected; cluster centre of mass: x = −3, y = −50, z = 30, mni_152 coordinates). (b) Mean grey matter density in the pCC cluster showing a significant difference between the matched groups (solid bars); distribution of values for all participants (triangles) overlaid with the first and third quartiles (box), interquartile range (whiskers), and density curve for each group. (c) Across all older adults (n = 93), second language proficiency is negatively correlated with grey matter density in the pCC (*r* = −0.30; *p* = 0.002, 2-tailed). pCC, posterior cingulate cortex; DMN, default mode network; *z*, standardized scores; * *p* < 0.05; ** *p* < 0.01.

### Monolingual older adults show greater default-executive coupling, with stronger RSFC between a FPCN hub node and the DMN

To test the hypothesis that bilingual older adults will show preserved intrinsic functional network integrity, despite lower neuroanatomical integrity, we used RSFC analysis to investigate within- and between-network connectivity among the nodes of three foundational large-scale networks – the FPCN, DMN, and DAN. The locations of 43 network nodes across these 3 networks were based on previous work that identified and characterized these networks based on both task-related and resting-state fMRI data (Spreng et al., 2013). The full matrices of the group means for all pairwise correlations between nodes for the monolingual and bilingual groups were calculated, and the latter subtracted from the former to produce the group difference matrix (Supplementary Fig. 2). The upper half of the matrix shows the group difference in mean pairwise correlation values; the lower half of the matrix is thresholded to show only the statistically significant differences (permutation test: q < 0.05, FDR-corrected for multiple comparisons across the entire matrix). Significant differences in RSFC between the monolingual and bilingual groups were highly specific and consistent with our hypotheses, with the monolinguals showing stronger RSFC between the aMFG in the left prefrontal cortex (PFC) and multiple nodes of the DMN only, including dorsomedial PFC, pCC, left posterior inferior parietal cortex, left superior frontal gyrus, left superior temporal sulcus, and ventromedial PFC (Fig. 2a,b). Mean RSFC between the aMFG and each of the 6 DMN nodes was also negatively correlated with second language proficiency across the entire sample of older adults (r = −0.29; p = 0.005, 2-tailed; Fig. 2c).

**Figure 2.**
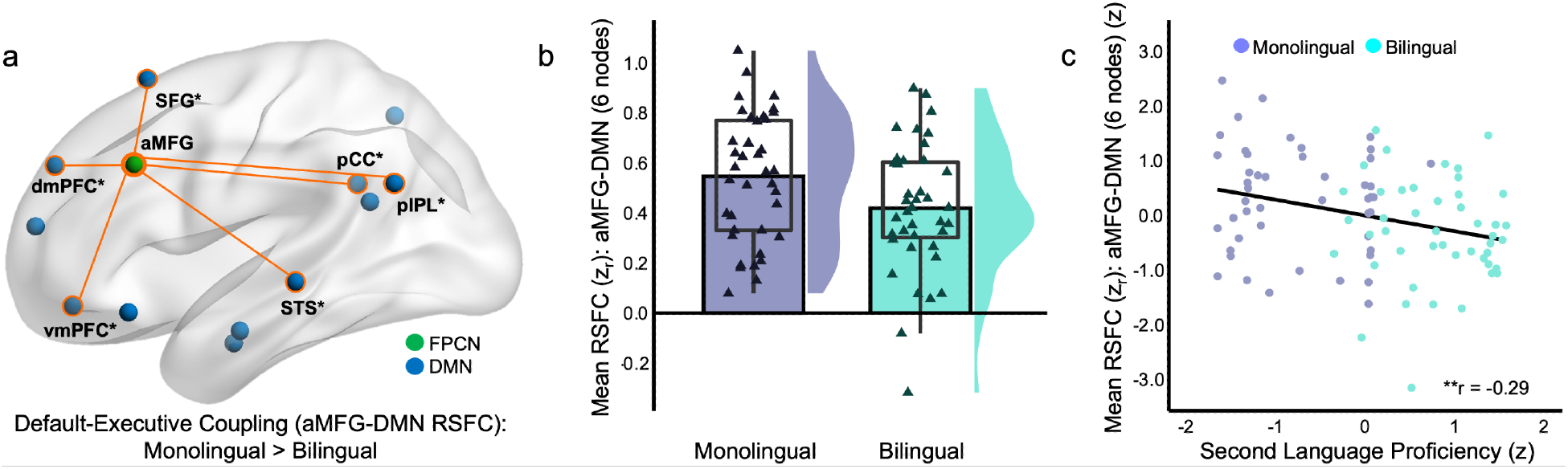
Differences in functional integrity estimated using RSFC. (a) Monolingual older adults show stronger RSFC of the left aMFG with multiple nodes of the DMN only (i.e., default-executive coupling), relative to bilinguals, including the dmPFC, pCC, left IPL, left SFG, left STS, and vmPFC (*p* < 0.05 FDR corrected for all pairwise correlations among all 43 nodes of the FPCN, DMN, and DAN) displayed on a glass brain representation of left hemisphere (see also Supplementary Fig. 2); orange lines/circles indicate significantly stronger pairwise RSFC in the monolingual older adults. (b) Mean RSFC between the aMFG and each of the 6 DMN nodes showing a significant difference between the matched groups (solid bars); distribution of values for all participants (triangles) overlaid with the first and third quartiles (box), interquartile range (whiskers), and density curve for each group. (c) Across all older adults (n=93), second language proficiency is negatively correlated with mean RSFC between the aMFG and the 6 nodes of the DMN (*r* = −0.29; *p* = 0.005, 2-tailed). FPCN, frontoparietal control network; DMN, default mode network; RSFC, resting-state functional connectivity; *zr*, Fisher *z*-transformed correlation; *z*, standardized score; for full list of all node label definitions, see Supplementary Table 1. * *p* < 0.05; ** *p* < 0.01.

Note that this pattern of increased functional connectivity between the MFG and DMN (i.e., default-executive coupling) is a hallmark characteristic of neurocognitive aging; it is associated with an inability to disengage the DMN during externally focussed tasks [task-related fMRI (Spreng and Schacter, 2012; Turner and Spreng, 2015)] and an increased reliance on semantic memory in the context of episodic memory decline (Spreng and Turner, 2019).

To further explore the extent of increased RSFC of the aMFG with the DMN, we used graph theory analysis to quantify the clustering coefficient of the aMFG with all nodes of DMN across the entire brain (see Fig. 3a for illustration of clustering calclulation, displayed in the left hemisphere). The monolingual group showed significantly higher clustering of the aMFG with the whole DMN than the bilingual group (mean difference = 0.20; permutation test: p = 0.04, 2-tailed) (Fig. 3b). Furthermore, clustering of the aMFG with the DMN was negatively correlated with second language proficiency across the entire sample of older adults (r = −0.22; p = 0.03, 2-tailed; Fig. 3c).

**Figure 3.**
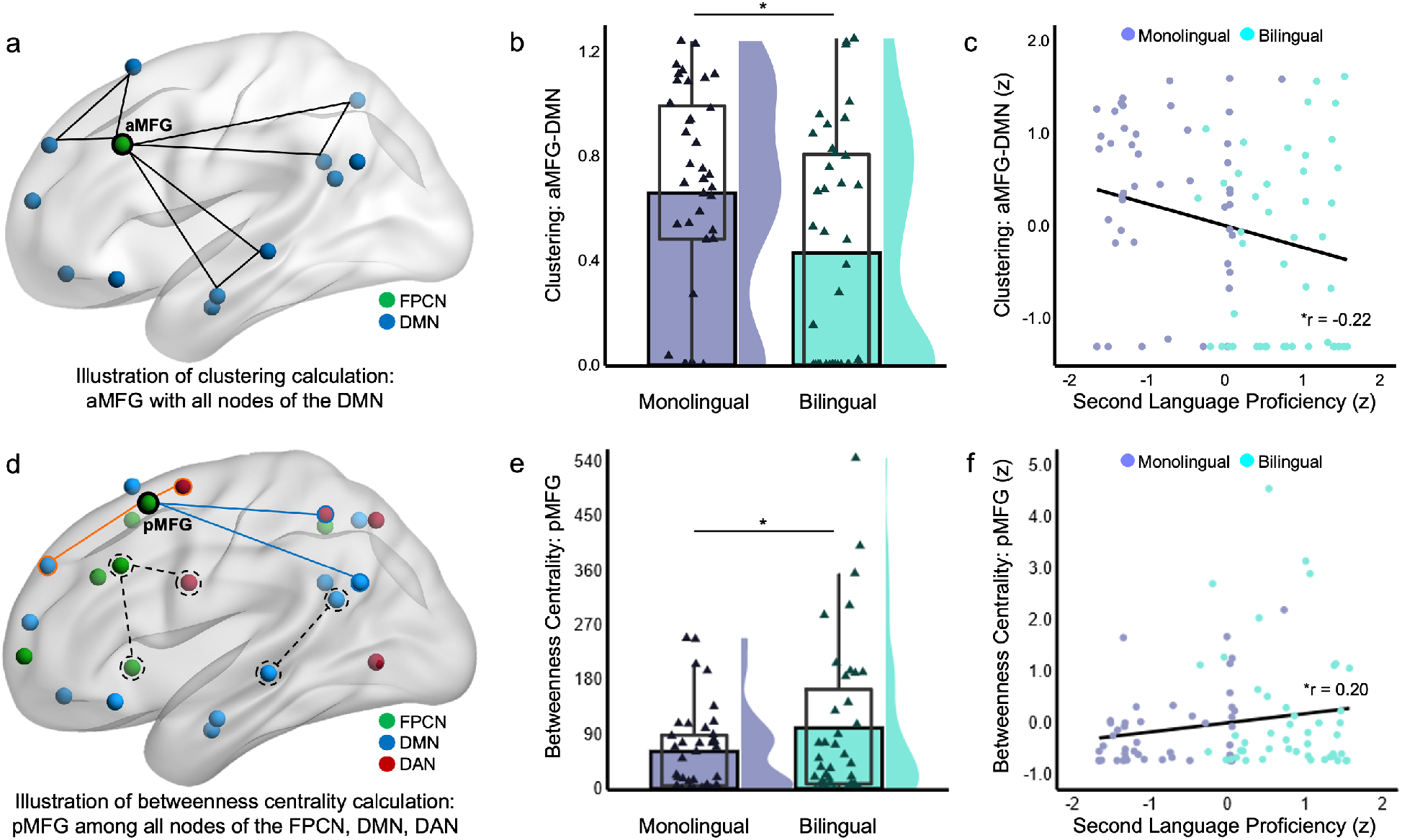
Differences in functional integrity estimated using graph theory analyses of RSCF. (a) Illustration of clustering coefficient calculation: number of complete triangles between the aMFG, a provincial hub of the FPCN, and every unique pair of nodes in the DMN, where triangle-sides (black lines) indicate statistically significant RSFC between nodes at a given density threshold. Illustration shows hypothetical clustering at a given threshold, displayed on a glass brain representation of the left hemisphere. (b) Monolingual older adults had higher clustering of the left aMFG with the DMN than bilinguals (mean clustering coefficient difference = 0.20; permutation test: *p* = 0.04, 2-tailed). Plot shows mean clustering coefficient (area under the curve) for each of the matched groups (solid bars); distribution of values for all participants (triangles) overlaid with the first and third quartiles (box), interquartile range (whiskers), and density curve for each group. (c) Across all older adults (n = 93), second language proficiency is negatively correlated with clustering of the aMFG with the DMN (*r* = −0.22; *p* = 0.03, 2-tailed). (d) Illustration of betweenness centrality calculation: number of shortest pairwise paths for all possible node-pairs among the 43 nodes of the FPCN, DMN, and DAN that pass through the left pMFG, a connector node of the FPCN. Illustration shows two shortest pairwise paths (solid lines) for 2 node-pairs (circled in orange and blue) that pass through the pMFG, and two shortest pairwise paths that do not (dashed black lines/circles). (e) Monolingual older adults had lower betweenness centrality of the left pMFG than monolinguals (mean difference = −45.06; permutation test: *p* = 0.04, 2-tailed). Plot shows mean betweenness centrality for each of the matched groups (solid bars); distribution of values for all participants (triangles) overlaid with the first and third quartiles (box), interquartile range (whiskers), and density curve for each group. (c) Across all older adults (n = 93), second language proficiency is positively correlated with betweenness centrality of the pMFG (*r* = 0.20; *p* = 0.04, 2-tailed). aMFG, anterior middle frontal gyrus; pMFG, posterior middle frontal gyrus; FPCN, frontoparietal control network; DMN, default mode network; DAN, dorsal attention network; RSFC, resting-state functional connectivity; *z*, standardized score. \**p* < 0.05.

### Bilingual older adults have higher betweenness centrality of a connector node of the FPCN

The FPCN facilitates control over one’s locus of attention by flexibly alternating its coupling between the DMN and DAN during internally vs. externally focussed cognition, respectively (Spreng et al., 2010). Previous work identified a key connector node of the FPCN in the pMFG with very high betweenness centrality, a graph theory analysis metric that quantifies the extent to which a node contributes to cross-network communication; thus, this node is thought to play a critical role in cognitive flexibility (Spreng et al., 2013). To assess potential differences in the flexibility of cross-network communication in monolingual vs. bilingual older adults, we compared the betweenness centrality of the pMFG (see Fig. 3d for illustration of betweenness centrality calculation, displayed in the left hemishpere) between the two groups. Bilingual older adults showed higher betweenness centrality of the pMFG than the monolinguals (mean difference = −45.06; permutation test: p = 0.04, 2-tailed; Fig. 3e). Finally, betweenness centrality of the pMFG was positively correlated with second language proficiency across the entire sample of older adults (r = 0.20; p = 0.04, 2-tailed; Fig. 3f).

## Discussion

Here, we demonstrate that when equated on cognitive performance using a rigorous matching procedure, bilingual older adults had lower grey matter density than their monolingual peers in the pCC, a hub of the DMN, which is particularly susceptible to structural and functional decline in aging and neurodegenerative disease (e.g., Alzheimer’s disease) (Buckner, 2005; Grady et al., 2006; Lustig et al., 2003). This finding confirms that lifelong bilingualism is a CR factor, as it demonstrates that bilingual older adults perform just as well as their monolingual peers on tasks of executive function despite showing signs of more advanced neuroanatomical aging. In contrast, the bilingual older adults had preserved intrinsic functional network structure, related to executive function and cognitive flexibility specifically, more consistent with that of younger brains (Spreng et al., 2013) than their monolinguals peers. While the monolingual older adults showed more prominent coupling of a hub node (aMFG) of the FPCN with the DMN, a hallmark characteristic of advanced age associated with a shift from fluid to crystalized cognition (Spreng et al., 2018), the bilinguals showed higher betweenness centrality of a connector node of the FPCN (pMFG), thought to underly cognitive flexibility (Spreng et al., 2013). Moreover, across all older adults, second language proficiency was negatively correlated with structural integrity, but positively correlated with multiple measures of intrinsic functional network integrity. These findings confirm that lifelong bilingualism, and second language proficiency in particular, contribute to CR, and support our hypotheses that this occurs through a process of bolstering the functional integrity, and maintaining neurocognitive flexibility, of the FPCN. Thus, a neural mechanism underlying CR in bilingualism is the experience-dependent maintenance of optimal intrinsic functional network structure in the face of neuroanatomical changes across the lifespan.

Several studies have investigated the possibility that lifelong bilingualism could lead to the preservation of, and/or compensatory increases in, the brain’s neuroanatomical structure by investigating white matter integrity using diffusion tensor imaging MRI. While some studies reported higher white matter integrity associated with bilingualism in older adults (John A.E. Anderson et al., 2018; Luk et al., 2011; Marin-Marin et al., 2020; Olsen et al., 2015), others reported prominent widespread declines in white matter integrity (Anderson et al., 2021; Brian T. Gold et al., 2013; Schweizer et al., 2012). A lack of consensus on the potential role of experience-dependent plasticity of brain structure in CR underscores the need for more research in this area using rigorous matching techniques to compare monolingual vs. bilingual groups – i.e., to delineate neuroanatomical differences associated with bilingualism per se, groups must be equated on cognitive performance (Anderson et al., 2021; John A. E. Anderson et al., 2018). One study that did this provided evidence of a more nuanced relationship between lifelong bilingualism and brain structural integrity in older adults. Cognitively healthy bilingual older adults had lower structural integrity of both grey and white matter globally, relative to monolinguals, but the bilinguals also had higher white and grey matter integrity within focal regions putatively involved in executive cognitive function specifically (Anderson et al., 2021).

Research has also investigated the possibility that differences in the way brain regions are engaged during task performance between monolingual and bilingual older adults could underlie or be a consequence of CR. Monolingual older adults show increased engagement of frontal brain regions or circuits reflective of increased cognitive effort at lower levels of task difficulty than bilingual adults, akin to differences between older and younger adults, respectively (Grundy et al., 2017). For example, monolingual older adults, despite having higher overall grey and white matter integrity than bilinguals, performed worse on a 2-back working memory task, and showed increased engagement of a frontostriatal circuit, including the left MFG in particular, during task performance (Anderson et al., 2021). Decreased task-related activation of frontal brain regions in bilingual older adults has been interpreted as increased efficiency (Grundy et al., 2017). However, the underlying structural or functional properties that could facilitate this more efficient task-related processing under certain task conditions have not been determined. We propose that intrinsic functional network connectivity – which is bolstered by task-related activity over time, which in turn facilitates future task performance – underlies the more efficient dynamic neurocognitive processing in bilingual older adults measured at the task-level.

Previous studies using fMRI to investigate potential differences in RSFC associated with bilingualism (Berken et al., 2016; Dash et al., 2022; DeLuca et al., 2019; Grady et al., 2015; Gullifer et al., 2018; Li et al., 2015; Luk et al., 2011; Marin-Marin et al., 2021; Thieba et al., 2019) have typically focussed on younger adults only, had small sample sizes, and/or failed to explicitly or adequately account for cognitive performance or neuroanatomical differences across groups of older individuals, complicating interpretations and limiting inferences about how observed changes are associated with CR per se. Nevertheless, our results are broadly consistent with previous work reporting increased RSFC primarily among regions involved in cognitive control (Grady et al., 2015) and attention networks specifically (Dash et al., 2022), in older bilingual adults. Previous work has also investigated differences in RSFC associated with various other CR factors across the lifespan (Anthony and Lin, 2018; Benson et al., 2018; Chan et al., 2021, 2018; N. Franzmeier et al., 2017; Nicolai Franzmeier et al., 2017a; Pietzuch et al., 2019; Stern et al., 2021; Weiler et al., 2018). One study demonstrated that level of education, as a proxy for CR, was positively correlated with increased RSFC between a left PFC region of the FPCN and the DAN, and negatively correlated with RSFC of this region with the DMN, in older adults with mild cognitive impairment (Nicolai Franzmeier et al., 2017b). The latter result parallels our findings of a negative correlation between second language proficiency and RSFC of the FPCN (aMFG) with the DMN, and increased RSFC between these regions in monolingual older adults, relative to bilinguals. This suggests that our discovery of the critical role of left MFG connectivity with other networks in CR associated with bilingualism might generalize to other CR factors as well, a hypothesis that should be investigated in future work using the analyses described here.

How could bilingualism have a long-term influence on RSFC of intrinsic functional networks? Managing multiple simultaneously active languages (Kroll et al., 2012; van Heuven et al., 2008) requires dynamically alternating the locus of one’s attention between the external environment, to monitor the situationally relevant language, and the content of one’s internal mental environment. Age-related memory declines are due, in part, to distraction resulting from a reduced ability to ignore task-irrelevant information (Healey et al., 2008; Stevens et al., 2008). Task-related modulation of functional connectivity of lateral PFC is critical for suppressing task-irrelevant processing in the brain (Zanto et al., 2011), which is reduced in older adults compared to young adults (Gazzaley et al., 2005). Moreover, task-related functional connectivity of the left lateral PFC with DMN regions specifically has been associated with memory failure in older adults due to an inability to ignore distraction (Stevens et al., 2008). The latter is consistent with research demonstrating that, unlike young adults, older adults “fail to deactivate” the DMN during externally focussed cognition (Grady et al., 2006; Lustig et al., 2003; Miller et al., 2008), and that this failure is associated with persistent functional connectivity of the DMN with the FPCN (Spreng and Schacter, 2012; Spreng and Turner, 2019; Turner and Spreng, 2015). Taken together, these results suggest that lateral PFC regions involved in cognitive control play a role in age-related differences in selective attention due to 1) reduced dynamic flexibility of functional connectivity with other brain regions/networks generally, and 2) persistent functional connectivity with the DMN, specifically. These task-related differences in older adults mirror age-related differences in intrinsic RSFC (Grady et al., 2016; Spreng et al., 2018), demonstrated most prominently by the monolingual older adults in this study, who showed lower cross-network RSFC of a connector node of the FPCN, and stronger RSFC of a hub node of the FPCN with the DMN. Previous work demonstrated that age-related differences in functional connectivity of the FPCN with other networks during both task and rest predicted cognitive performance (Grady et al., 2016). Thus, the differences reported here between monolingual and bilingual older adults in cross-network RSFC of two key nodes of the FPCN (aMFG and pMFG) could account for the preservation of cognitive function in bilinguals in the face of more advanced neuroanatomical aging, relative to monolinguals.

Younger looking patterns of RSFC in the bilingual older adults could be the consequence of a long history of increased demand on the FPCN to facilitate attentional control. Metaphorically, just as targeted physical exercise builds specific muscles of the body needed to optimize physical performance over time, “mental exercise” may strengthen RSFC among specific brain regions needed to optimize cognitive performance. Experience-dependent changes in RSFC are highly specific to the brain regions engaged during task performance (Albert et al., 2009; Lewis et al., 2009; Stevens et al., 2010; Tambini et al., 2010). Further, the magnitude of experience-dependent changes in RSFC predicts the degree of improvement in future task performance (Lewis et al., 2009; Stevens et al., 2010; Tambini et al., 2010). Thus, the pattern of preserved RSFC of the FPCN hub and connector nodes in the brains of bilingual older adults is highly consistent with the pattern of task-evoked activity that facilitates domain-general attentional control, needed to cognitively manage multiple languages simultaneously.

Recent work sought to identify a RSFC connectome associated with cognitive reserve using a data-driven approach, which could be useful as a direct measure of cognitive reserve and a potential outcome measure for interventions aimed at improving CR (Stern et al., 2021). The pattern of RSFC identified was correlated with IQ, and notably, included primarily frontal brain regions. Indeed, understanding the factors that give rise to CR and its underlying neural mechanism(s) is an important endeavour, with broad implications ranging from the level of the individual to society. For example, a recent large-scale longitudinal study demonstrated relationships between educational attainment, RSFC of the brain’s functional networks, and neurocognitive health in older age (Chan et al., 2021). In general, older adults show reduced within-network RSFC and increased between-network RSFC relative to young adults, reflecting a widespread pattern of dedifferentiation [also referred to as reduced segregation (Wig, 2017)] of functional network organization broadly (Chan et al., 2014; Geerligs et al., 2015; Grady et al., 2016; Setton et al., 2022; Spreng et al., 2016). This effect of aging is more prominent in older adults without a college degree than their college-educated peers, and predicts impending progression of dementia up to a decade later (Chan et al., 2021).

Our results are generally consistent with previous work described above reporting large-scale network dedifferentiation in aging and widespread, somewhat non-specific differences in patterns of RSFC associated with CR (e.g., education and IQ). Importantly, however, our results further delineate the specific processes that lead to network-wide changes at the level of individual network nodes, revealing their dissociable roles in maintaining within- and between-network communication in the context of a well-established CR factor – bilingualism. Notably, the pattern of cross-network functional connectivity of MFG regions that was more prominent in bilingual older adults and correlated with second language proficiency in the current study is the reverse of the pattern associated with the well-established shift from fluid to crystalized cognition in typical aging (Spreng and Turner, 2019). Therefore, our findings highlight the critical roles of specific brain regions – i.e., discrete nodes of the FPCN within the MFG – in maintaining domain-general attentional control in aging, and suggest the intriguing possibility that the pattern of RSFC of these regions associated with bilingualism could be a neural mechanism underlying multiple CR factors. Understanding these factors is critical for informing personal lifestyle choices and policy decisions in an increasingly older global population.

## Materials and Methods

### Participants

Ninety-three participants (age: mean ± SD = 74.04 ± 3.86, range = 68-85; 63 female) were recruited from the local area surrounding Toronto, Ontario, Canada; each participant was a resident of Canada at the time of testing. All participants were right-handed and had no history of heart disease, neurological or psychological disorders, or traumatic brain injuries. Data from a subset of these participants were analyzed in previous work (Anderson et al., 2021; John A.E. Anderson et al., 2018). An additional 3 participants were recruited but to ensure data quality, these participants were excluded from analyses due to head motion in the MRI scanner exceeding 3mm in one or both of the resting-state fMRI runs.

The Language and Social Background Questionnaire [LSBQ (John A. E. Anderson et al., 2018)] was used to categorize participants as monolingual or bilingual. Participants were interviewed via telephone to validate bilingual status, and those who could not be reliably identified as either monolingual or bilingual were not included in the study. Of the 93 participants analyzed, 47 participants (29 female) were identified as bilingual and 46 (34 female) were identified as monolinguals. All participants completed the Mini-Mental State Examination [MMSE (Folstein et al., 1975)] and D-KEFS battery (Delis et al., 2001). The D-KEFS is a well-established battery that assesses a comprehensive variety of cognitive processes related to executive control.

### Propensity Score Matching

Interpreting group differences in brain structure and function related to cognitive reserve requires that monolingual and bilingual groups have an equivalent level of cognitive functioning. Therefore, in the current study, monolinguals and bilinguals were matched on multiple neuropsychological and demographic variables using propensity-score matching (PSM). A propensity score for each individual was obtained by using logistic regression to predict group membership given a set of observed covariates. Participants from one group were then matched to those in the second group based on the propensity scores (Rosenbaum and Rubin, 1983). Nearest neighbour matching was used to match the propensity scores of individuals between groups. PSM accounts for multivariate interaction, minimizes selection error when using a large number of matching variables, and is well-suited for smaller sample sizes that are usually found in neuroimaging studies (Austin and Steyerberg, 2015; Ho et al., 2007; Rosenbaum and Rubin, 1983).

To prioritize matching the monolingual and bilingual groups on cognitive performance, PSM was done using the MatchIt R package (Ho et al., 2011) in a 2-stage hierarchical manner. The first level matched the groups on 4 neuropsychological measures: MMSE, Trail Making Task (letter-number-switching score), Shipley-2 verbal score, and Shipley-2 block patterns score Using the propensity scores, each bilingual participant was matched to a monolingual; bilinguals that could not be matched, and any remaining monolinguals, were removed from the matched-group analyses. Once the groups were matched on cognitive performance, they were then matched on 3 demographic variables using the same procedure: age, education, and sex. The final group sizes were 39 monolinguals (age: mean ± SD = 73.51 ± 3.25, range = 68-79; 30 female) and 39 bilinguals (age: mean ± SD = 73.87 ± 4.00, range = 69-83; 23 female).

### MRI Data Acquisition

MRI data were collected in Toronto, Ontario, Canada at York University using a Siemens Trio 3T scanner with a 32-channel head coil. For each participant, a high-resolution T1-weighted anatomical scan was acquired (TR = 1.9 s, TE = 2.52 ms, 192 axial slices, slice thickness = 1 mm, field of view (FOV) = 256 mm, 256 x 256 acquisition matrix). After the anatomical scan, participants completed two six-minute, multi-echo “resting-state” functional scans, wherein participants were instructed to remain still in the scanner while visual maintaining fixation on a cross (TR = 3 s, TE= 14 ms, 30 ms, 46 ms, flip angle = 83°, 43 axial slices, FOV = 216 mm, voxel size = 3 mm isotropic).

### Voxel-Based Morphometry

To assess differences in grey matter integrity, voxel-based morphometry (VBM) was conducted using the FSL-VBM software suite (Douaud et al., 2007; Smith et al., 2004). T1-weighted structural images were brain-extracted (BET) to remove skull. Next, tissue-type segmentation was conducted (FAST4) to quantify grey matter density at the voxel-wise level across the whole brain for each participant. Grey matter images were normalized to a study-specific grey matter template through a two-step process. First, all grey matter volume images were non-linearly aligned to the MNI_152 grey matter template and averaged to create a study specific grey matter template; second, the native-space grey matter image for each participant was non-linearly aligned to the study specific template (FNIRT). The registered grey matter images were corrected for local expansion or contraction (caused by the non-linear component of the spatial transformation) by dividing by the Jacobian of the warp field. The modulated images were then smoothed with an isotropic Gaussian kernel of 3 mm full width at half maximum (FWHM). Group differences in grey matter density were determined using voxel-wise *t*-tests (3dttest++) in Analysis of Functional Neuroimages [AFNI (Cox, 1996)] with alpha < 0.05; correction for multiple comparisons was done using the AFNI non-parametric equitable thresholding and clustering (ETAC) method (Cox, 2019), with 10 voxel-wise *p*-thresholds ranging from *p* = 0.01 to *p* = 0.001. To calculate the correlation between second language proficiency and grey matter density in the pCC across all participants (n = 93), a single significant cluster from the group analysis was used as a ROI; mean grey matter probability across all voxels within this ROI was calculated for each participant.

### fMRI Data Preprocessing

Echo-planar images were preprocessed using the AFNI multi-echo independent component analysis (ME-ICA) processing pipeline (DuPre et al., 2021; Kundu et al., 2013, 2012). For each echo of both resting-state runs, any large transient spikes of activity were identified and removed via interpolation (3dDespike). Data were then slice-time corrected (3dTshift) and aligned to the initial volume (3dVolreg). ME-ICA (tedana) was conducted to further remove noise components related to motion and physiological artifacts (Kundu et al., 2013, 2012). The combination of multi-echo fMRI and ME-ICA has been shown to be more effective at removing spurious noise sources than conventional methods (Power et al., 2018). This denoising technique leverages the TE-dependence of the transverse relaxation rate, R_2_*; almost all fMRI signal, regardless of origin, can be expressed as some combination of R_2_*, and initial signal intensity, S_0_ (Wu and Li, 2005). All echoes are combined and analyzed using ICA to determine the degree to which independent components were associated with R_2_* changes. Components highly associated with R_2_* mechanisms were identified as BOLD signal components, whereas weakly associated components were identified as noise and removed (Kundu et al., 2013, 2012). The effectiveness of ME-ICA denoising obviates the need for global signal regression (Spreng et al., 2019). Data for each run were then smoothed with an isotropic Gaussian kernel of 6 mm FWHM (3dBlurInMask) and mean-scaled. Finally, the 2 scaled runs were concatenated and transformed into a study specific MNI_152-template space (@auto_tlrc).

### Resting-State Functional Connectivity

Coordinates for all nodes of the DMN, FPC, and DAN were obtained from a previous study that identified and characterized these network-nodes based on both task-related and resting-state fMRI data; nodes were assigned network labels corresponding to their “rest network affiliation” (Spreng et al., 2013). For each participant, nodes were defined by spherical regions of interest (ROIs) with a radius of 6 mm centered on the *a priori* coordinates. To quantify resting-state functional connectivity (RSFC), the mean BOLD signal time-series across all voxels within each node was extracted and pairwise Pearson correlation coefficients were calculated for all node-pairs and transformed to z-values using Fisher’s r-to-z transformation. This yielded a 43 by 43 node-wise correlation matrix for each participant. Significant differences in RSFC between the monolingual and bilingual groups were determined using permutation tests (1000 permutations) for every node-pair and false discovery rate correction (threshold: q < 0.05, FDR corrected) for multiple comparisons across the entire matrix.

### Graph Theoretic Analysis

For each participant, a graph was constructed from the node-wise correlation matrix using the GraphVar software suite (Kruschwitz et al., 2015). To account for potential threshold effects, we calculated graph metrics across a series of network densities (density range = 0.22-0.4). Two graph metrics were calculated and reported here: weighted local clustering coefficient and betweenness centrality. The weighted clustering coefficient of a node quantifies how tightly clustered a given node is with its immediate neighbours, while considering the weight of the connections; it is calculated as the sum of the weights of all closed triplets centered on a node at a given density, normalized by the sum of the weights of all possible triplets centered around that node. The betweenness centrality of a node is the number of shortest paths between all possible node-pairs that pass through the given node at a given density; betweenness centrality is a measure of the extent to which a node contributes to communication across networks, and thus can be used to identify main interconnector nodes (Rubinov and Sporns, 2010). Both metrics were normalized by calculating the mean area under the curve across the range of densities. Significant group differences were determined using permutation tests (1000 permutations).

### Brain-Behaviour Correlations

Previous behavioural research has implicated the active use and proficiency of a second language as the critical factor having a positive effect on cognitive reserve (Calabria et al., 2020). Second language proficiency was previously quantified from the LSBQ. A factor analysis on all the LSBQ items identified a latent variable with clustered items representing second language proficiency. The eigenvalue from this factor was used as a subscale of the LSBQ to quantify the active use and proficiency of a second language (John A. E. Anderson et al., 2018). To determine if second language proficiency is related to structural and functional brain properties, we used Pearson correlation to quantify the relationship between second language proficiency and each of our brain measures across the entire group of 93 participants (age: mean ± SD = 74.21 ± 4.09, range = 68-85; 65 female). Second language proficiency was correlated with each of the following brain measures: PCC grey matter density, mean RSFC between the aMFG and the 6 DMN nodes identified in the group comparison of the full correlation matrix (dmPFC, pCC, left IPL, left SFG, left STS, and vmPFC), clustering coefficient of the aMFG with the entire DMN, and betweenness centrality of the pMFG. The significance of the Pearson correlation was determined using two-tailed tests at alpha < 0.05.

## Supplementary Material

**SupplementaryFigure 1:**
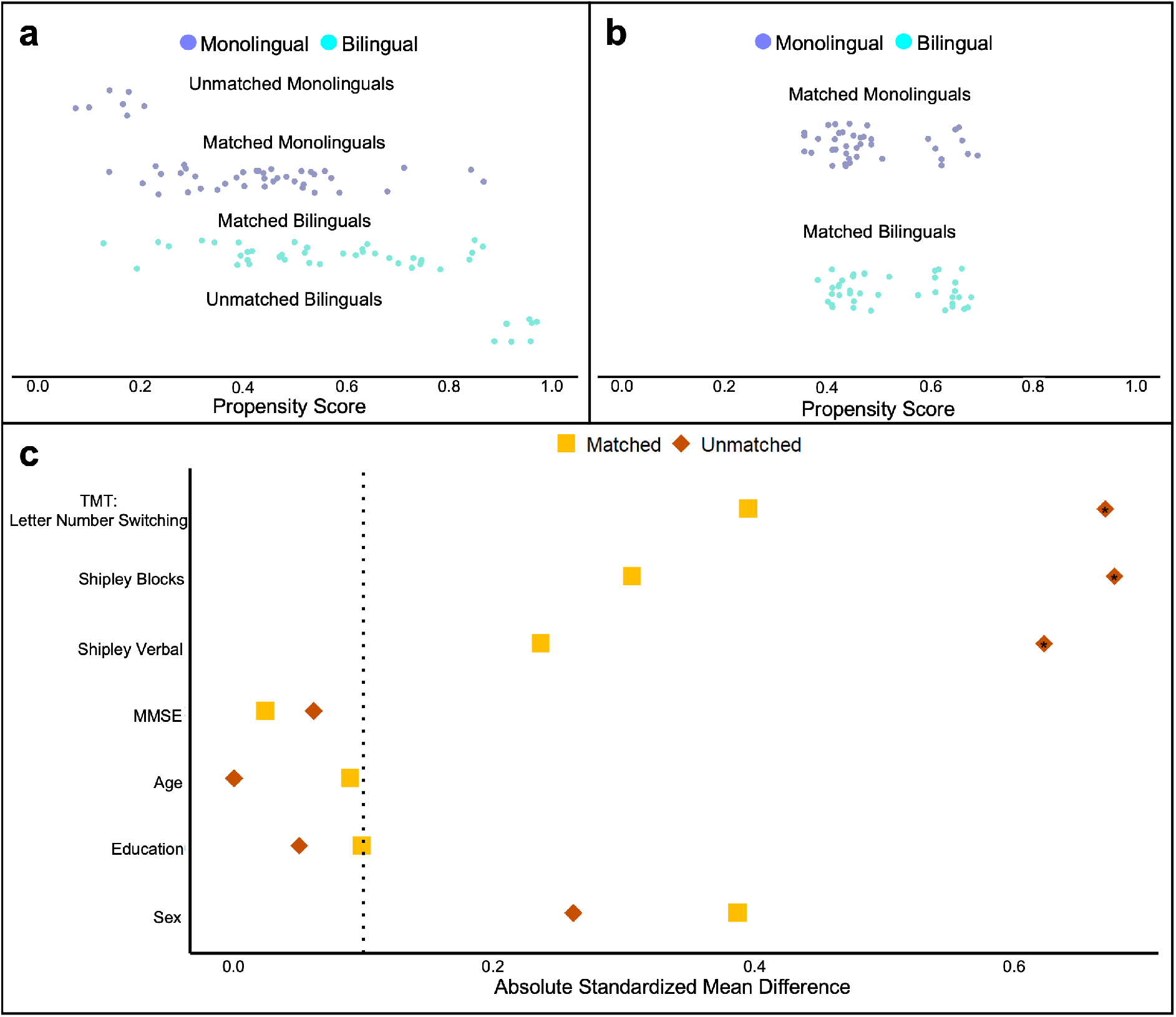
Propensity score matching (PSM) of monolingual and bilingual groups. (**a**) Distribution of propensity scores for the matched monolingual (n = 39) and bilingual (n = 39) groups and excluded (unmatched) participants following PSM on cognitive measures. (**b**) Distribution of propensity scores for the matched monolingual and bilingual groups following PSM on demographic variables, which did not change group membership. (**c)** Absolute standardized mean difference between monolinguals and bilinguals on each of the cognitive and demographic variables for the matched and unmatched groups. Significant differences (indicated by an asterisk) between the unmatched monolingual and bilingual groups on three cognitive measures (TMT Letter Number Switching, Shipley Blocks, and Shipley Verbal scores) were eliminated in the matched groups following PSM. TMT, Trail Making Task; MMSE, Mini * p < 0.01

**SupplementaryFigure 2:**
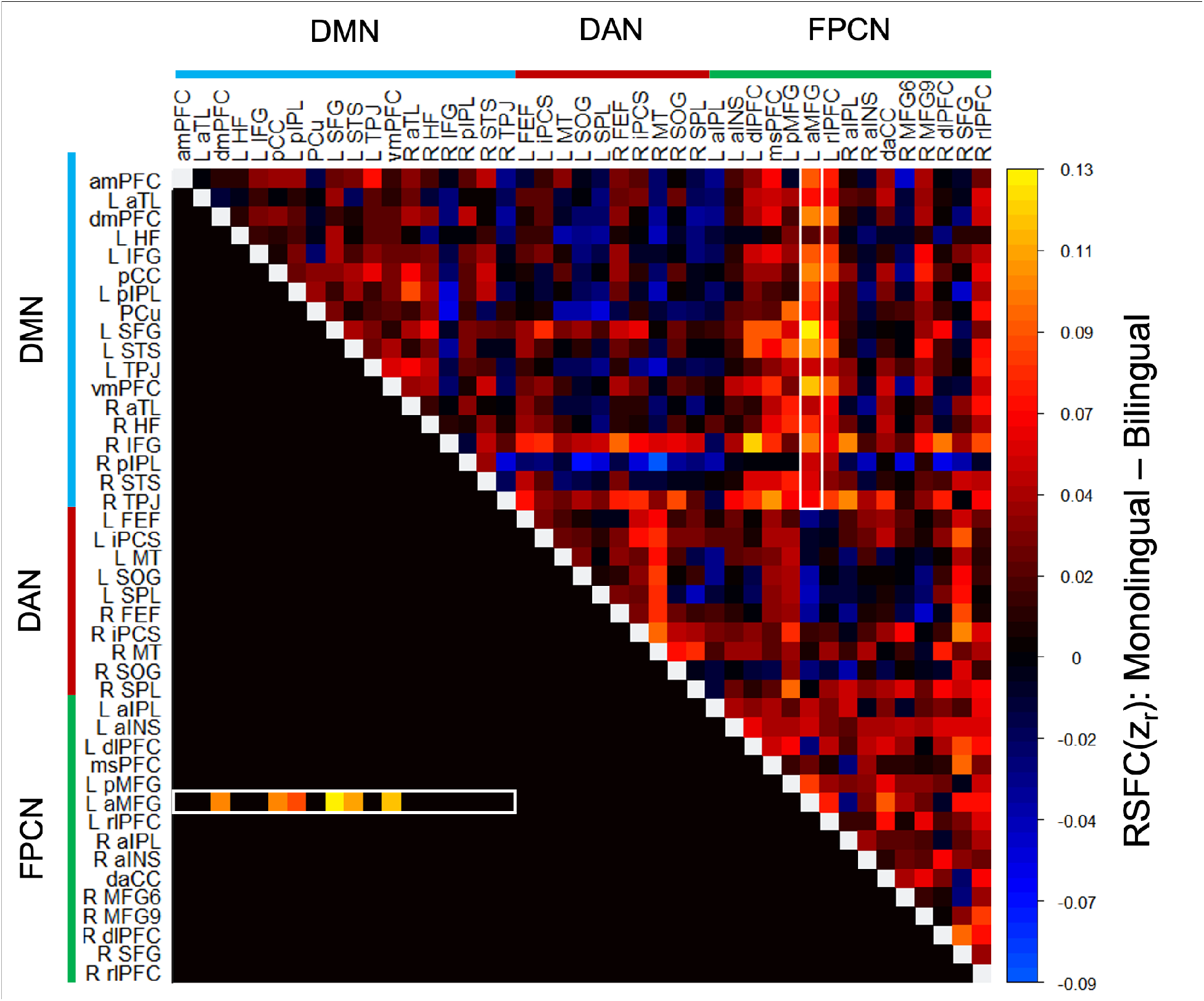
Differences in RSFC within and between the DMN, DAN, and FPCN. Monolingual older adults show stronger RSFC of the left aMFG with multiple nodes of the DMN, relative to bilinguals. Full pairwise correlation matrix of all 43 nodes of the FPCN, DMN, and DAN: upper half of the matrix shows group differences (monolingual - bilingual) in mean RSFC for all pairwise correlations between nodes; lower half of the matrix shows only the statistically significant differences (permutation test: q < 0.05, FDR corrected). The left aMFG showed stronger RSFC with 6 nodes of the DMN: dmPFC*, pCC*, left IPL*, left SFG**, left STS*, and vmPFC*. RSFC, resting-state functional connectivity; zr, Fisher z-transformed correlation; FPCN, frontoparietal control network; DMN, default mode network; DAN, dorsal attention network; for full list of all node label definitions, see Supplementary Table 1. * p < 0.002; ** p < 0.001

**Supplementary Table 1.**
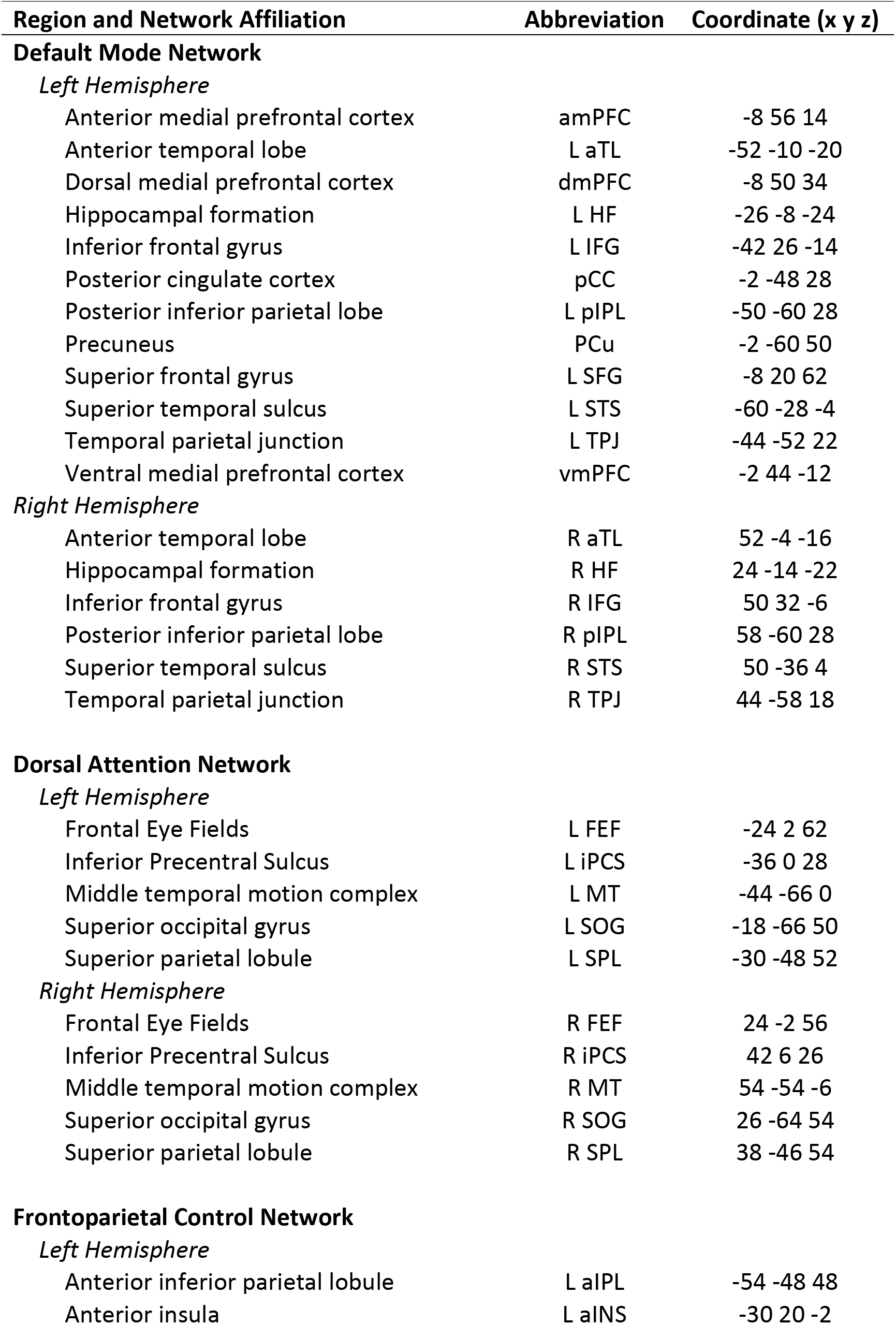
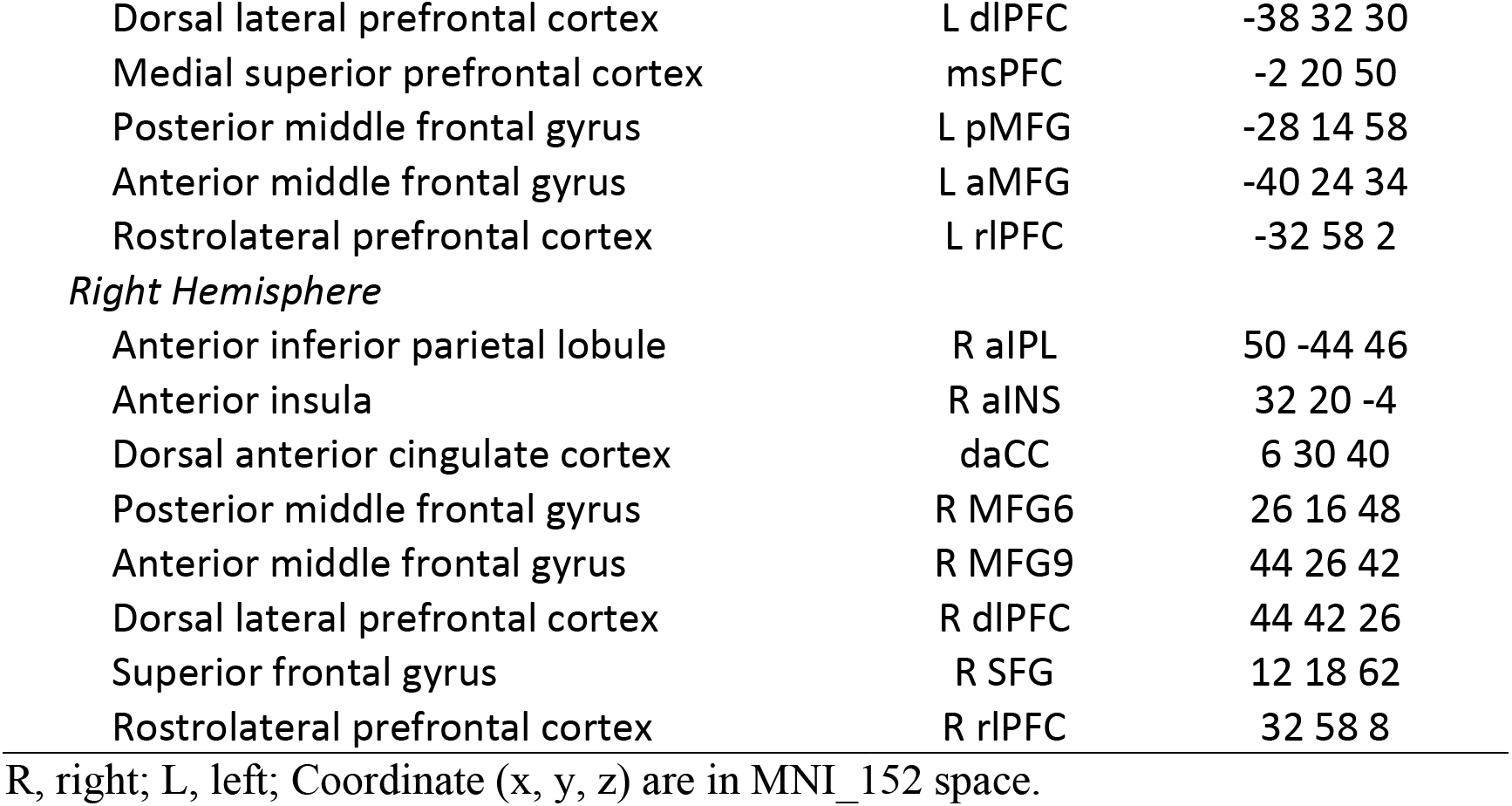
List of all nodes of default mode network, dorsal attention network, and frontoparietal control network, and their abbreviations.

