## Supplementary Material for "A neural mechanism of cognitive reserve: The case of bilingualism"

**Affiliations:**
[1]Department of Psychology, York University, Toronto, Canada.
[2]Department of Cognitive Science, Carleton University, Ottawa, Canada.
[3]Rotman Research Institute at Baycrest Hospital, Toronto, Canada.
[4]Departments of Psychology and Psychiatry, University of Toronto, Toronto, Canada.

### Supplementary Material

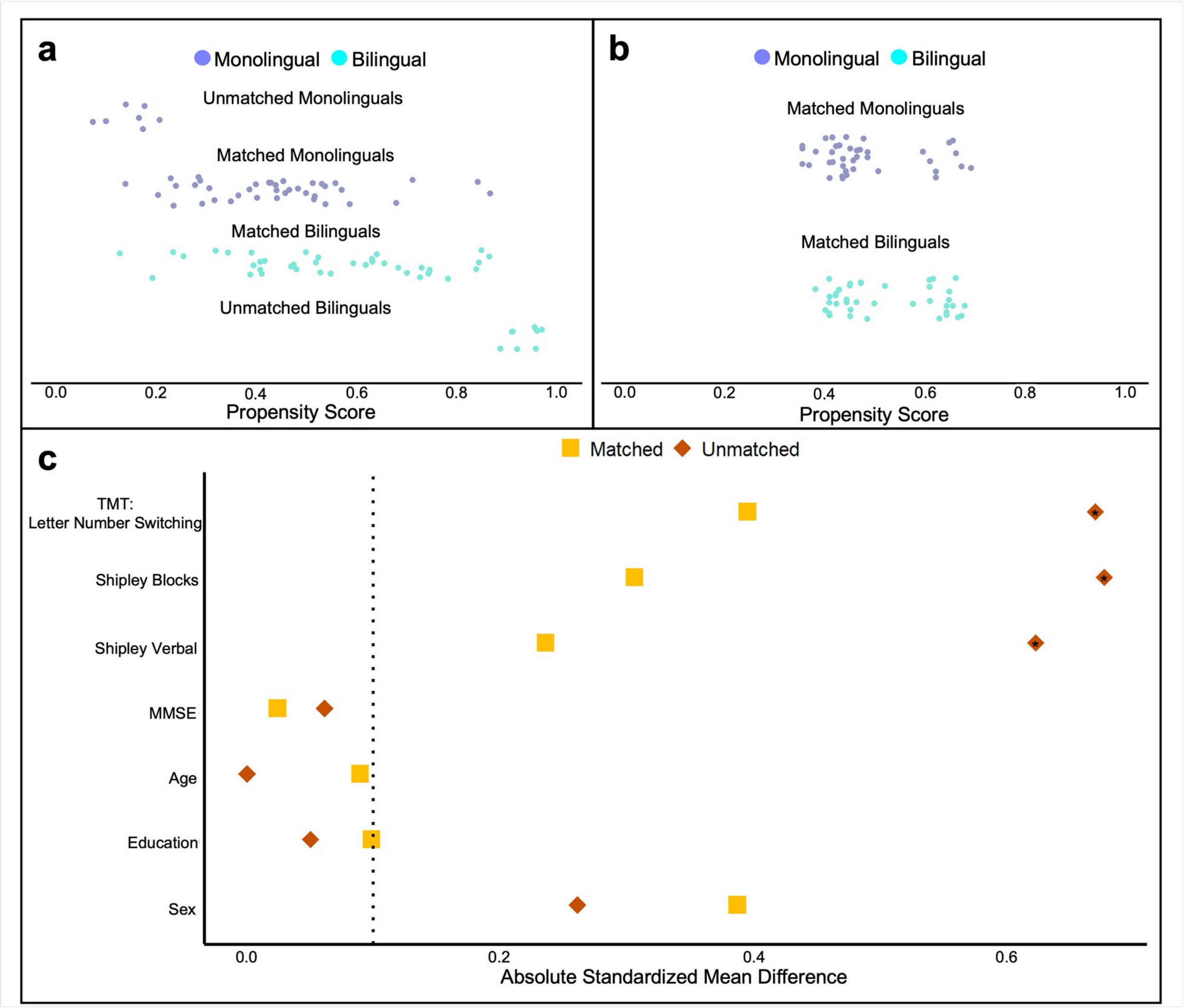

**SupplementaryFigure 1:** Propensity score matching (PSM) of monolingual and bilingual groups. (**a**) Distribution of propensity scores for the matched monolingual (n = 39) and bilingual (n = 39) groups and excluded (unmatched) participants following PSM on cognitive measures. (**b**) Distribution of propensity scores for the matched monolingual and bilingual groups following PSM on demographic variables, which did not change group membership. (**c)** Absolute standardized mean difference between monolinguals and bilinguals on each of the cognitive and demographic variables for the matched and unmatched groups. Significant differences (indicated by an asterisk) between the unmatched monolingual and bilingual groups on three cognitive measures (TMT Letter Number Switching, Shipley Blocks, and Shipley Verbal scores) were eliminated in the matched groups following PSM. TMT, Trail Making Task; MMSE, Mini * $p < 0.01$

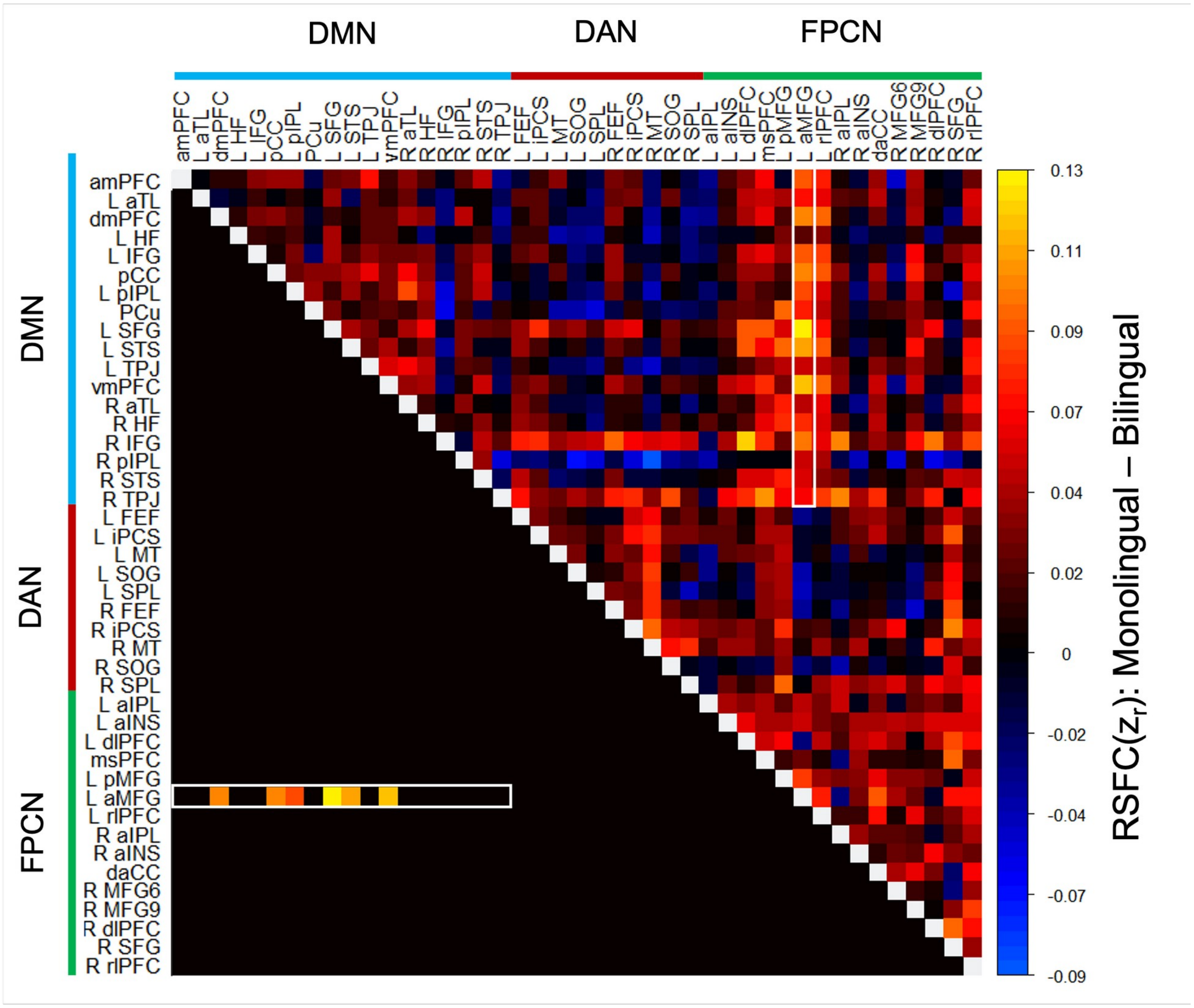

**Supplementary Figure 2:** Differences in RSFC within and between the DMN, DAN, and FPCN. Monolingual older adults show stronger RSFC of the left aMFG with multiple nodes of the DMN, relative to bilinguals. Full pairwise correlation matrix of all 43 nodes of the FPCN, DMN, and DAN: upper half of the matrix shows group differences (monolingual - bilingual) in mean RSFC for all pairwise correlations between nodes; lower half of the matrix shows only the statistically significant differences (permutation test: $q < 0.05$, FDR corrected). The left aMFG showed stronger RSFC with 6 nodes of the DMN: dmPFC*, pCC*, left IPL*, left SFG**, left STS*, and vmPFC*. RSFC, resting-state functional connectivity; $z_r$, Fisher z-transformed correlation; FPCN, frontoparietal control network; DMN, default mode network; DAN, dorsal attention network; for full list of all node label definitions, see Supplementary Table 1. * $p < 0.002$; ** $p < 0.001$

Supplementary Table 1. List of all nodes of default mode network, dorsal attention network, and frontoparietal control network, and their abbreviations.

| Region and Network Affiliation | Abbreviation | Coordinate (x y z) |
|---|---|---|
| **Default Mode Network** | | |
| *Left Hemisphere* | | |
| Anterior medial prefrontal cortex | amPFC | -8 56 14 |
| Anterior temporal lobe | L aTL | -52 -10 -20 |
| Dorsal medial prefrontal cortex | dmPFC | -8 50 34 |
| Hippocampal formation | L HF | -26 -8 -24 |
| Inferior frontal gyrus | L IFG | -42 26 -14 |
| Posterior cingulate cortex | pCC | -2 -48 28 |
| Posterior inferior parietal lobe | L pIPL | -50 -60 28 |
| Precuneus | PCu | -2 -60 50 |
| Superior frontal gyrus | L SFG | -8 20 62 |
| Superior temporal sulcus | L STS | -60 -28 -4 |
| Temporal parietal junction | L TPJ | -44 -52 22 |
| Ventral medial prefrontal cortex | vmPFC | -2 44 -12 |
| *Right Hemisphere* | | |
| Anterior temporal lobe | R aTL | 52 -4 -16 |
| Hippocampal formation | R HF | 24 -14 -22 |
| Inferior frontal gyrus | R IFG | 50 32 -6 |
| Posterior inferior parietal lobe | R pIPL | 58 -60 28 |
| Superior temporal sulcus | R STS | 50 -36 4 |
| Temporal parietal junction | R TPJ | 44 -58 18 |
| **Dorsal Attention Network** | | |
| *Left Hemisphere* | | |
| Frontal Eye Fields | L FEF | -24 2 62 |
| Inferior Precentral Sulcus | L iPCS | -36 0 28 |
| Middle temporal motion complex | L MT | -44 -66 0 |
| Superior occipital gyrus | L SOG | -18 -66 50 |
| Superior parietal lobule | L SPL | -30 -48 52 |
| *Right Hemisphere* | | |
| Frontal Eye Fields | R FEF | 24 -2 56 |
| Inferior Precentral Sulcus | R iPCS | 42 6 26 |
| Middle temporal motion complex | R MT | 54 -54 -6 |
| Superior occipital gyrus | R SOG | 26 -64 54 |
| Superior parietal lobule | R SPL | 38 -46 54 |
| **Frontoparietal Control Network** | | |
| *Left Hemisphere* | | |
| Anterior inferior parietal lobule | L aIPL | -54 -48 48 |

| | | |
|---|---|---|
| Anterior insula | L aINS | -30 20 -2 |
| Dorsal lateral prefrontal cortex | L dlPFC | -38 32 30 |
| Medial superior prefrontal cortex | msPFC | -2 20 50 |
| Posterior middle frontal gyrus | L pMFG | -28 14 58 |
| Anterior middle frontal gyrus | L aMFG | -40 24 34 |
| Rostrolateral prefrontal cortex | L rlPFC | -32 58 2 |
| *Right Hemisphere* | | |
| Anterior inferior parietal lobule | R aIPL | 50 -44 46 |
| Anterior insula | R aINS | 32 20 -4 |
| Dorsal anterior cingulate cortex | daCC | 6 30 40 |
| Posterior middle frontal gyrus | R MFG6 | 26 16 48 |
| Anterior middle frontal gyrus | R MFG9 | 44 26 42 |
| Dorsal lateral prefrontal cortex | R dlPFC | 44 42 26 |
| Superior frontal gyrus | R SFG | 12 18 62 |
| Rostrolateral prefrontal cortex | R rlPFC | 32 58 8 |

R, right; L, left; Coordinate (x, y, z) are in MNI_152 space.
